# Oligo-barcodes illuminate holocentric karyotype evolution in *Rhynchospora* (Cyperaceae)

**DOI:** 10.1101/2023.10.31.564898

**Authors:** Yennifer Mata-Sucre, Leticia Maria Parteka, Christiane Ritz, Andrés Gatica-Arias, Leonardo P. Félix, Willian Wayt Thomas, Gustavo Souza, André L. L. Vanzela, Andrea Pedrosa-Harand, André Marques

## Abstract

Holocentric karyotypes are assumed to rapidly evolve through chromosome fusions and fissions due to the diffuse nature of their centromeres. Here, we took advantage of the recent availability of a chromosome-scale reference genome for *Rhynchospora breviuscula*, a model species of this holocentric genus, and developed the first set of oligo-based barcode probes for a holocentric plant. These probes were applied to 13 additional species of the genus, aiming to investigate the evolutionary dynamics driving the karyotype evolution in *Rhynchospora*. The two sets of probes were composed of 27,392 (green) and 23,968 (magenta) oligonucleotides, and generated 15 distinct FISH signals as a unique barcode pattern for the identification of all five chromosome pairs of the *R. breviuscula* karyotype. Oligo-FISH comparative analyzes revealed different types of rearrangements, such as fusions, fissions, putative inversions and translocations, as well as genomic duplications among the analyzed species. Two rounds of whole genome duplication (WGD) were demonstrated in *R. pubera*, but both analyzed accessions differed in the complex chain of events that gave rise to its large, structurally diploidized karyotypes with 2*n* = 10 or 12. Considering the phylogenetic relationships and divergence time of the species, the specificity and synteny of the probes were maintained up to species with a divergence time of ∼25 My. However, karyotype divergence in more distant species hindered chromosome mapping and the inference of specific events. This barcoding system is a powerful tool to study chromosomal variations and genomic evolution in holocentric chromosomes of *Rhynchospora* species.

## INTRODUCTION

Despite their essentiality for proper chromosome segregation, the organization of centromeres is diverse among eukaryotes (Drinnenberg and Akiyoshi, 2017). Based on their chromosomal localization, two main configurations are recognized: monocentromeres and holocentromeres, referring to the localized, size-restricted or the diffuse distribution of centromeric activity along the chromosome, respectively (Melters *et al.,* 2012; Wong *et al.,* 2020; Schubert *et al.,* 2020). The peculiar diffuse structural organization of the latter allows spindle fibers to bind along the entire length of the so-called holocentric chromosomes. Thus, fragments originated from chromosomal breaks and fusions can be inherited during cell division more frequently compared to their monocentric counterparts (Vanzela and Colaço 2002; Melters *et al.,* 2012; Heckmann and Houben 2013; Jankowska *et al.,* 2015). In species with holocentric chromosomes, fusions and fissions do not appear to disrupt proper segregation, so holocentricity has been assumed to potentially reduce or eliminate selective pressure against chromosomal rearrangements, triggering reproductive isolation, i.e., chromosomal speciation (Lucek *et al.,* 2022).

Holocentromeres have arisen several times independently among animals and plants, such as in the Cyperaceae (sedges) family (Melters *et al.,* 2012; Escudero *et al.,* 2016; Senaratne *et al.,* 2022). The genus *Rhynchospora* Vahl (beaksedges) is the third largest clade of Cyperaceae (Araújo *et al.,* 2012), with about 400 species distributed worldwide (POWO, 2023). Chromosomal numbers range from 2*n* = 4 in *R. tenuis* Link (Vanzela 1996) to 2*n* = 61 in *R. globosa* (Kunth) Roem. & Schult. (Burchardt *et al*., 2020), with a most likely ancestral chromosomal number of *n* = 5 (Burchardt *et al.,* 2020; Hofstatter *et al.,* 2022). Models of chromosomal evolution point to polyploidy as the main driver in *Rhynchospora*, followed by dysploidy (Ribeiro *et al.,* 2018; Burchardt *et al.,* 2020; Hofstatter *et al.,* 2022). Recently, we described an auto-octoploid origin for *Rhynchospora pubera* (Vahl) Boeckeler (2*n* = 10), formed after two rounds of genome duplication. Remarkably, post-polyploidy genome shuffling events due to end-to-end chromosome fusions substantially reduced its chromosome number to the ancestral *n* = 5, demonstrating that extensive chromosomal rearrangements underlie rapid karyotype evolution in the genus (Hofstatter *et al.,* 2022). Although chromosome fusions were independently observed in both *R. tenuis* (2*n* = 4) and *R. pubera* (2*n* = 10) (Hofstatter *et al*., 2022), chromosome number variation was only observed in *R. tenuis*, as depicted from its highly reduced karyotype. Thus, it is likely that chromosomal rearrangements may still be hidden in species having the common ancestral chromosomal number *n* = 5.

Whole genome comparison might be considered the most comprehensive way to shed light on chromosomal evolution. However, most plant groups still have only a single reference genome, making comparative genomic analyses between closely related species difficult and requiring the use of alternative feasible methods (Parween *et al.,* 2015; Qin *et al.,* 2019; Escudero *et al.,* 2023). Chromosome barcoding with oligo-based probes has been used to determine karyotypes and investigate chromosomal rearrangements, meiotic pairing, and recombination in a wide range of species (Braz *et al.,* 2018; Meng *et al.,* 2018; Li *et al.,* 2021; Do Vale Martins *et al.,* 2021; De Oliveira Bustamante *et al.,* 2021; Doležalová *et al.,* 2022; Nascimento and Pedrosa-Harand, 2023). Oligo-FISH barcoding is based on the production of two different oligomer libraries from multiple regions of multiple chromosomes to produce an unique barcode signal pattern for each individual chromosome pair, facilitating the unambiguous identification of all chromosomes of a species in a single FISH experiment (Braz *et al.,* 2018, 2020; Liu and Zhang 2021; De Oliveira Bustamante *et al.,* 2021). The availability of sequenced and assembled genomes for *R. pubera*, *R. tenuis* and *R. breviuscula* H. Pfeiff. (Hofstatter *et al.,* 2022) provide the unique opportunity to design oligo-FISH probes for the creation of a universal oligo-FISH barcoding system to study genomic evolution in the genus *Rhynchospora*. Here we developed an oligo-FISH barcoding system and performed a comparative genomic study in the holocentric genus *Rhynchospora*, comparing collinear regions to identify major chromosome structural changes among *Rhynchospora* species. We demonstrate that the oligo-FISH-based barcode technique is a powerful tool for chromosome identification and karyotype evolution research even in highly dynamic holocentric karyotypes.

## RESULTS

### Development of barcoding probes for chromosome identification in *Rhynchospora* species

We have developed two barcode-oligo probes (Rbv-I and Rbv-II) for chromosome identification in *Rhynchospora* species. These probes comprise 27,392 and 23,968 oligonucleotides (45-nt), respectively, which were identified in the five chromosomes of the *R. breviuscula* reference genome (Hofstatter *et al.,* 2022). The Rbv-I probe (green signals) covered 8 different regions, whereas the Rbv-II probe (magenta signals) covered 7 regions in the five *Rhynchospora* chromosomes (**Fig. 1a**). Each probe covered between 585 and 1,110 kb of a chromosomal segment in the pseudomolecule and was comprised around 3,424 oligonucleotides (**Table 1**). The oligo-FISH barcode libraries, hereafter referred to as oligo-probes, were designed to achieve a density between 4.13 and 4.19 oligos/kb in the region of interest to ensure good visibility of the hybridization signals after FISH (**Table 1, Table S1**).

**Figure 1:**
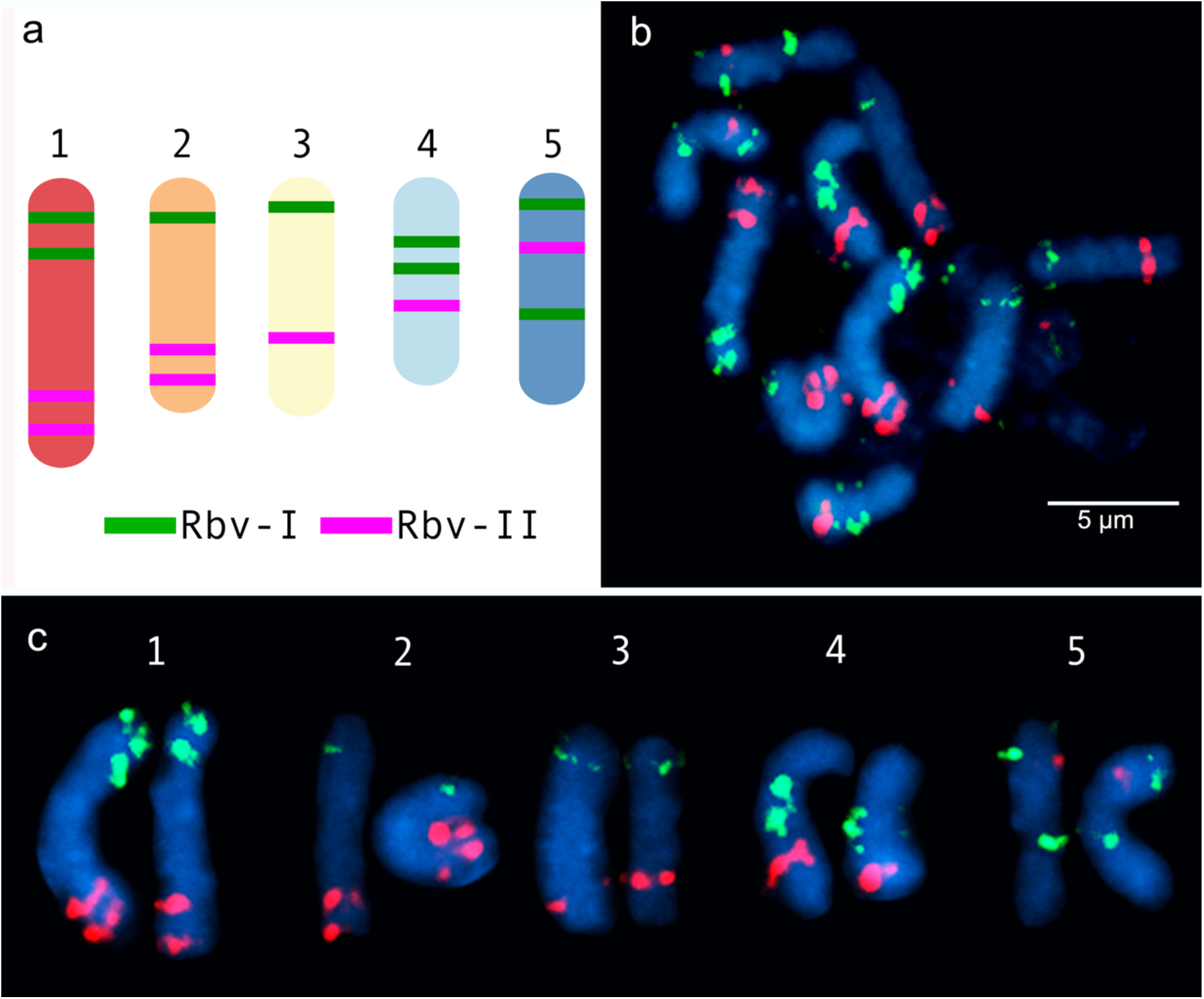
Predicted location and mapping of oligo-FISH barcode signals in *Rhynchospora breviuscula*. (**a**) Predicted oligos were selected from a total of 15 chromosomal regions (8 green "Rbv-I" and 7 magenta "Rbv-II" regions). The five chromosome pairs can be distinguished from each other based on the number and location of the Rbv-I/Rbv-II signals. (**b**) FISH mapping of metaphase mitotic chromosomes of *R. breviuscula* using oligo-barcode probes Rbv-I (green) and Rbv-II (magenta). (**c**) Karyogram identifying homologous chromosomes I to V from the same cells shown in **b** of *R. breviuscula*. Chromosomes were counterstained with DAPI (blue). Bar = 5 µm.

**Table 1.**
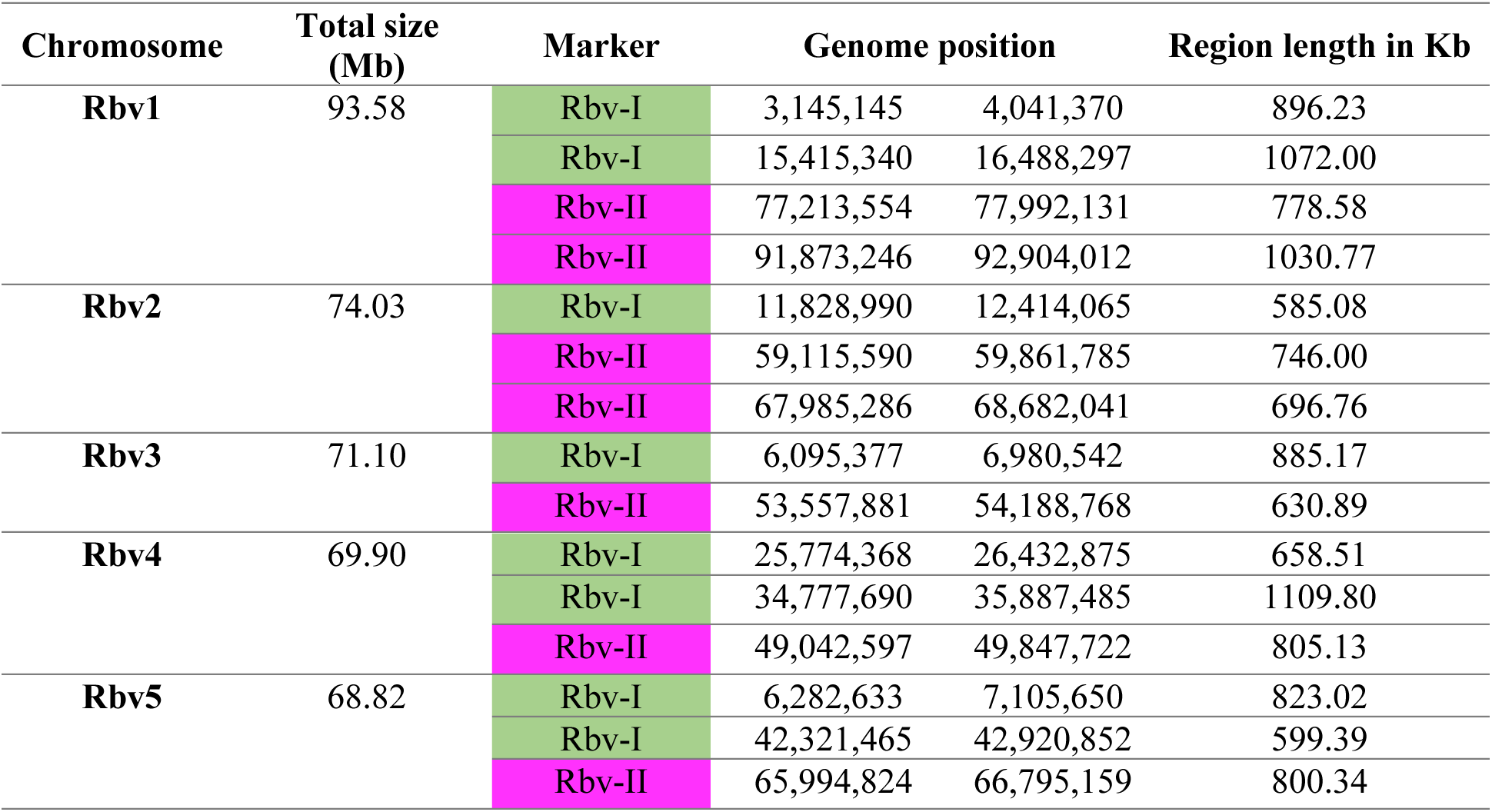
Genomic positions of oligo barcodes in the reference genome of *Rhynchospora breviuscula* (Rbv). Total size and region’s length were calculated using the available reference genome (Hofstatter *et al*., 2022). The color of each chromosome marker was defined in accordance with **Fig. 1** description.

| Chromosome | Total size (Mb) | Marker | Genome position |  | Region length in Kb |
| --- | --- | --- | --- | --- | --- |
| **Rbv1** | 93.58 | Rbv-I | 3,145,145 | 4,041,370 | 896.23 |
|  |  | Rbv-I | 15,415,340 | 16,488,297 | 1072.00 |
|  |  | Rbv-II | 77,213,554 | 77,992,131 | 778.58 |
|  |  | Rbv-II | 91,873,246 | 92,904,012 | 1030.77 |
| **Rbv2** | 74.03 | Rbv-I | 11,828,990 | 12,414,065 | 585.08 |
|  |  | Rbv-II | 59,115,590 | 59,861,785 | 746.00 |
|  |  | Rbv-II | 67,985,286 | 68,682,041 | 696.76 |
| **Rbv3** | 71.10 | Rbv-I | 6,095,377 | 6,980,542 | 885.17 |
|  |  | Rbv-II | 53,557,881 | 54,188,768 | 630.89 |
| **Rbv4** | 69.90 | Rbv-I | 25,774,368 | 26,432,875 | 658.51 |
|  |  | Rbv-I | 34,777,690 | 35,887,485 | 1109.80 |
|  |  | Rbv-II | 49,042,597 | 49,847,722 | 805.13 |
| **Rbv5** | 68.82 | Rbv-I | 6,282,633 | 7,105,650 | 823.02 |
|  |  | Rbv-I | 42,321,465 | 42,920,852 | 599.39 |
|  |  | Rbv-II | 65,994,824 | 66,795,159 | 800.34 |

To validate the accuracy of the Rbv-I and Rbv-II oligo-probes, hybridizations of mitotic metaphase cells were performed in the same *R. breviuscula* genotype used as the reference genome (Hofstatter *et al*., 2022). The green and magenta FISH probes generated bright and specific signals on all 10 homologous chromosomes, matching to the predicted patterns and validating the chromosome specificity of the oligo-probes (**Fig. 1b**). The positions of the signals formed a barcode pattern that uniquely identified the five chromosome pairs of *R. breviuscula* (**Fig. 1c**). The FISH signals of the Rbv-I probes (green) were weaker than those of the Rbv-II probe (**magenta, Fig. 1c**).

### Comparative karyotype evolution within *Rhynchospora* species revealed by barcode oligo-FISH

After establishing the karyotype identification system of *R. breviuscula*, we performed comparative oligo-FISH on 13 additional *Rhynchospora* species using the two probes developed here to reveal karyotype evolution in the genus (**Fig. 2 & Fig. S1**). Our sampling comprised species of all main five clades of the genus according to their phylogenetic relationships (Costa *et al.,* 2023), except for clade III. Karyotype analysis revealed holocentric chromosomes in all species analyzed, evident from the lack of primary constriction, with chromosome numbers ranging from 2*n* = 4 to 2*n* = 26 and chromosome sizes ranging from 1.10 µm (*R. alba*) to 16.43 µm (*R. pubera*; **Table S2**). The two oligo-probes generated recognizable barcode signals in most species of clade IV, similar to the observed for *R. breviuscula*. The observed patterns allowed us to infer multiple chromosome fusions, putative inversions and translocations in different chromosomes of some species, revealing that intra- and interchromosomal rearrangements occurred during the divergence of species from clade IV (**Fig. 2 & Fig. S1**). The barcode patterns were, however, less evident in species of clade I, II and V, congruent to the phylogenetic distances among clades, and limiting the overall use of this set of probes for a broader karyotype analysis. Nevertheless, we have outlined the main chromosome rearrangements in the genus below.

**Figure 2:**
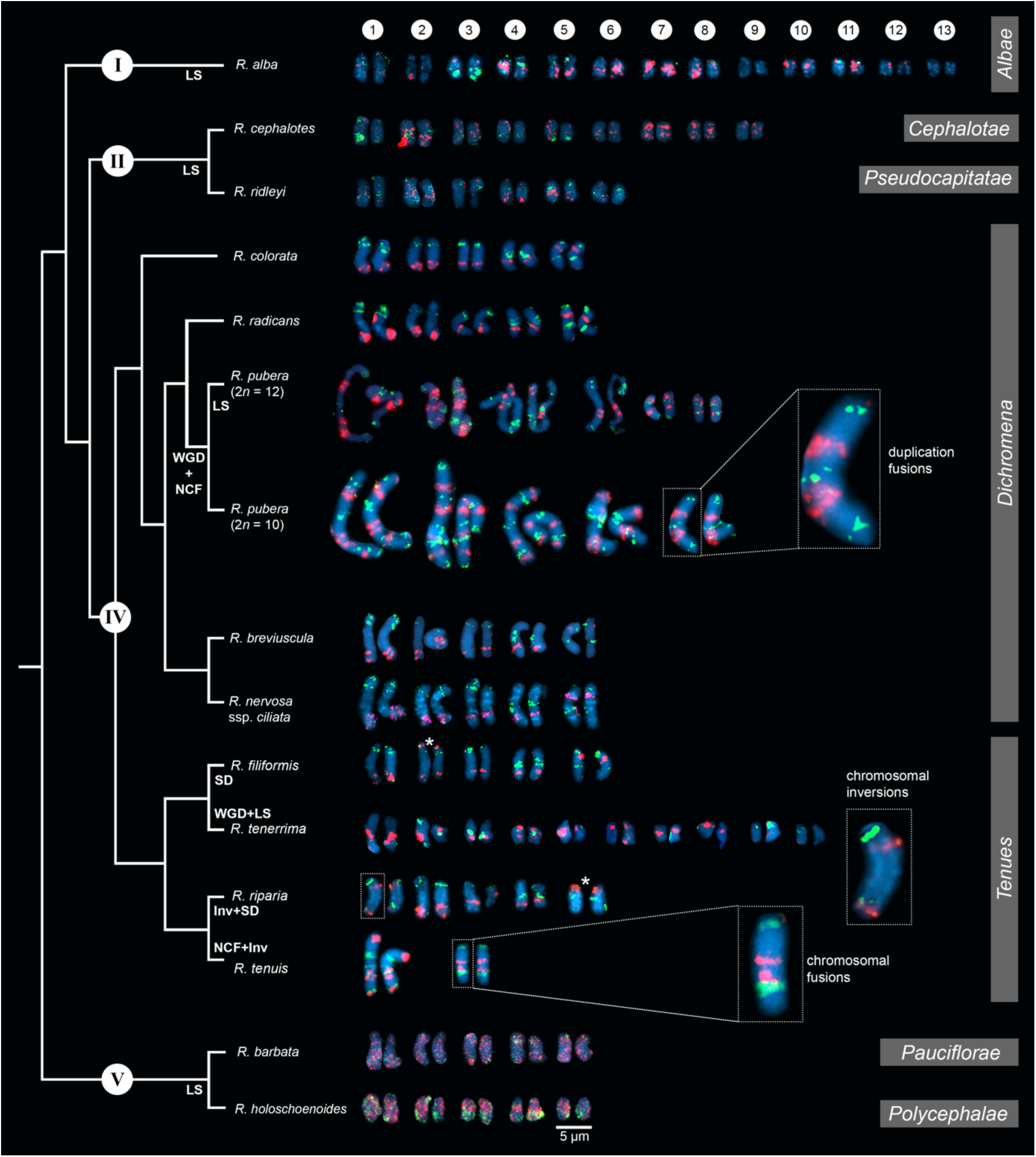
Comparative karyograms of 14 *Rhynchospora* species based on the barcode oligo-probes Rbv-I (green) and Rbv-II (magenta). For each species, chromosome pairs are ordered from left to right according to their putative *R. breviuscula* orthologs. Insets illustrate chromosome fusions in *R. pubera* and *R. tenuis*, and an inversion in chr1 in *R. riparia*. Asterisk points to signal duplication in chr2 and chr5 of *R. filiformis* and *R. riparia*, respectively, likely due to chromosomal translocations. For species without clear oligo-probe patterns, chromosome pairs were arranged according to their decreasing sizes. NCF indicates fusion of nested chromosomes; Inv refers to inversions, LS refers to loss of signal (deletions), while SD refers to signal duplications. WGD denotes whole genome duplication. Clade numbering and phylogenetic relationships are based on Costa *et al*. (2023). Section assignments (grey bars on the right) are given based on recent phylogenetic and taxonomical revisions (Thomas *et al.,* 2009; Buddenhagen *et al*., 2016; Silva Filho *et al.,* 2021). Bar = 5 µm.

#### *Clade IV – Section* Dichromena *(Michx.) Griseb*

The oligo-probes Rbv-I and Rbv-II were hybridized to the somatic chromosomes of five species of the section *Dichromena*, i.e., *R breviuscula*, *R. colorata* (L.) H. Pefeiff., *R. nervosa* (Vahl) Boeckeler ssp. *ciliata* (G. Mey.) T. Koyama, and *R. radicans* (Schltdl. & Cham.) H. Pfeiff.; as well as two accessions of *R. pubera*, our reference genotype with 2*n* = 10 (Hofstatter *et al*., 2022) and a recently collected accession with 2*n* = 12, referred to here as "*R. pubera*-10" and "*R. pubera*-12", respectively (**Fig. 2 & Fig. 3**). Green and magenta signals derived from the two probes confirmed our recent findings (Hofstatter *et al*., 2022) that the *R. pubera*-10 karyotype has originated from two rounds of whole genome duplication followed by a complex chain of end-to-end chromosome fusion events. Indeed, we observed a signal pattern congruent with four copies of each of the five chromosomes of *R. breviuscula* (**Fig. 2**, **Fig. 3a & Fig. S2**). The *R. pubera*-12 accession was initially thought to be a dysploid cytotype of *R. pubera*-10 due to a single fission of one chromosome pair. Nevertheless, our detailed analysis revealed a much more complex karyotype. We found this accession to have additional rearrangements besides the genome duplications and fusions found in *R. pubera*-10, as indicated from the FISH signal patterns observed from its six homologous chromosomes. The FISH pattern observed in *R. pubera*-12 suggests fission of one chromosome pair and loss of significant chromosomal regions likely due to massive genomic reshuffling (**Fig. 2 & Fig. 3b**). Although the quality of the FISH signals was similar to those of *R. breviuscula*, we could not find a pattern that allowed us to decipher its complex karyotype. Indeed, we could only count a total of 43 FISH signals, 21 of the Rbv-I probe and 22 of the Rbv-II probe, in contrast to the 60 signals observed in *R. pubera*-10 (**Fig. 3a–b**). Even though a resolution/condensation matter could be hiding additional signals, these findings are consistent with the loss of multiple genomic regions equivalent to a set of *R. breviuscula* chromosomes during the karyotype evolution of *R. pubera*-12 (**Fig. 2**).

**Figure 3:**
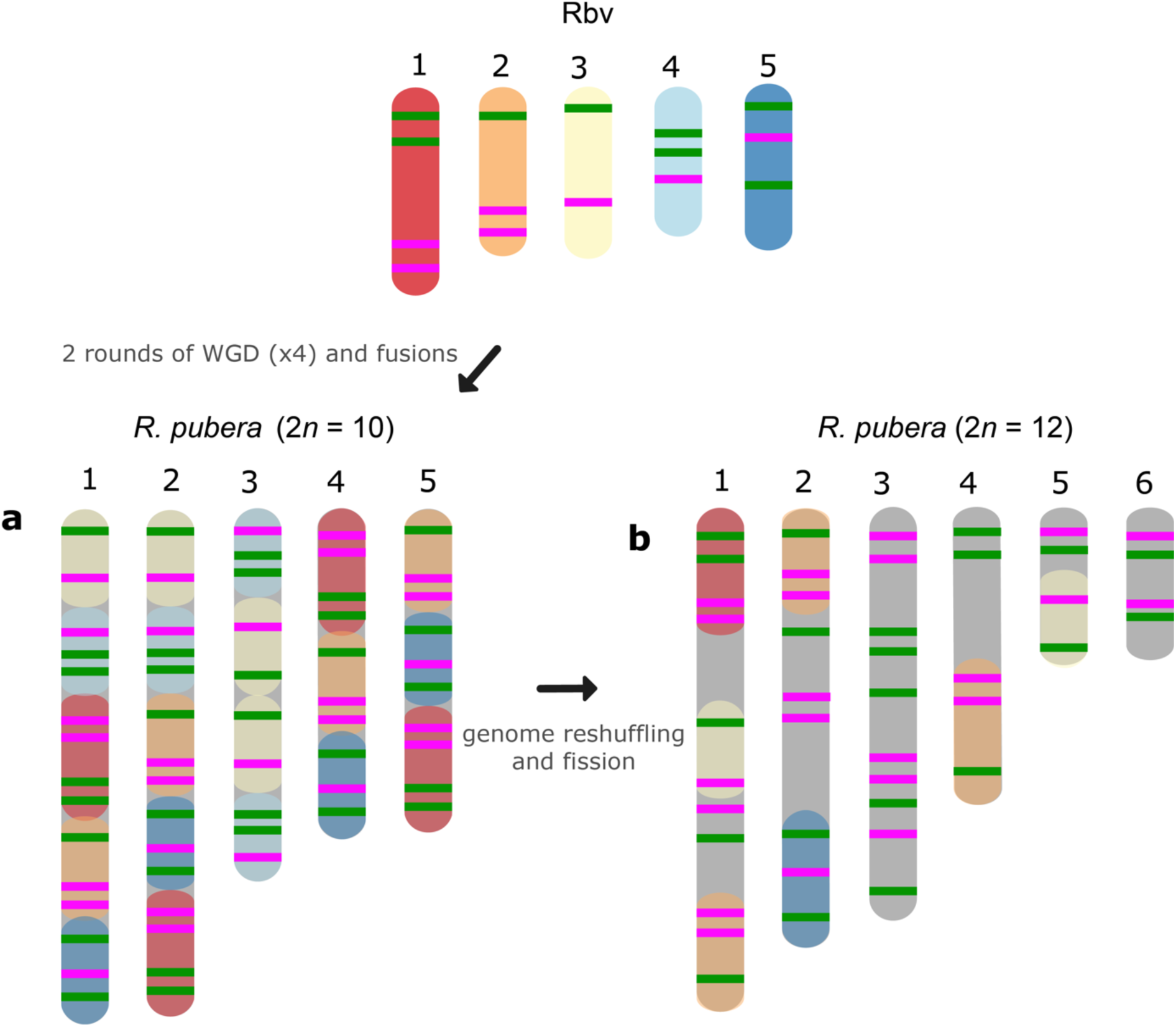
Schematic idiograms of the chromosome fusions identified in *R. pubera* using *R. breviuscula* (Rbv-I and Rbv-II) oligo-probes. (**a**) Fusions of four copies of the five chromosomes of *R. breviuscula* detectable in *R. pubera*-10. As expected, we found 8×4 green signals and 7×4 magenta signals totaling 60 signals in the haploid *R. pubera* genome. (**b**) Despite finding some similar patterns to *R. breviuscula*, we were unable to decipher the complete karyotype of *R. pubera* with 2*n* = 12. It is possible that there are additional complex chromosome rearrangements than what we observed in the sample with 2*n* = 10. Indeed, the reduced number of signals, 21 green and 22 magenta, suggests that this accession had undergone considerable genome downsizing. Fusions of these chromosomes are predicted based on oligo-FISH barcode modifications and chromosome sizes.

Hybridization signals of Rbv-I and Rbv-II oligoprobes on the chromosomes of *R. colorata* (2*n* = 10), *R. nervosa* ssp. *ciliata* (2*n* = 10), and *R. radicans* (2*n* = 10) matched exactly the same pattern as those of *R. breviuscula*, indicating a high synteny and collinearity among species with same ploidy and chromosome number in the section *Dichromena* (**Fig. 2 & Fig. S1**).

#### *Clade IV – Section* Tenues *Kük*

Species of this section, *R. tenuis, R. riparia* (Nees) Boeckeler, *R. filiformis* Vahl and *R. tenerrima* Nees ex Spreng., presented rearranged patterns of signals for Rbv-I and Rbv-II oligoprobes, showing fusions, inversions, translocations and duplications with respect to the *R. breviuscula* pattern (**Fig. 2 & 4**). A putative intrachromosomal translocation and a duplication was observed on chromosome 2 of *R. filiformis* (2*n* = 10; **Fig. 4b**), in which one of the magenta signals translocated closer to the opposite end of the chromosome adjacent to the green signal, which is duplicated, different from that observed in the *R. breviuscula* pattern. A large inversion on chromosome I of *R. riparia* (2*n* = 10) would explain the changes observed in the FISH signal pattern (**Fig. 4c**). Duplication and deletion of magenta or green oligo-barcode signals were also observed on chromosome 5 of *R. filiformis* and *R. riparia*, indicating additional small structural rearrangements in these species (**Fig. 4b-c**). *Rhynchospora tenuis* (2*n* = 4: **Fig. 4d**) showed all the FISH signal patterns of the five *R. breviuscula* chromosomes on its two chromosome pairs, except for a putative inversion on chromosome 2, confirming the fusion events of chromosomes 1-2-5 and 3-4 on its chromosomes 1 and 2, respectively (**Fig. 4d**), as we have recently shown by genome assembly (Hofstatter *et al*., 2022). Another interesting case is found in *R. tenerrima*, a species with 2*n* = 20, which is assumed to be a polyploid (Arguelho *et al*., 2012). The hybridization pattern of Rbv-I and Rbv-II FISH probes revealed 21 signals in total (11 green and 10 magenta, **Fig. 4e**), in contrast to the expected 15 for a diploid if compared to *R. breviuscula* chromosomes. Thus, it is possible for this species to be polyploid, but due to the reduced number of signals expected for a fully duplicated set of chromosomes, i.e., 30, *R. tenerrima* is likely already experiencing diploidization through genome downsizing. Indeed, this is evidenced from the lack of a similar barcode pattern to *R. breviuscula* chromosomes, which hampered us to decipher its karyotype (**Fig. 4e**).

**Figure 4:**
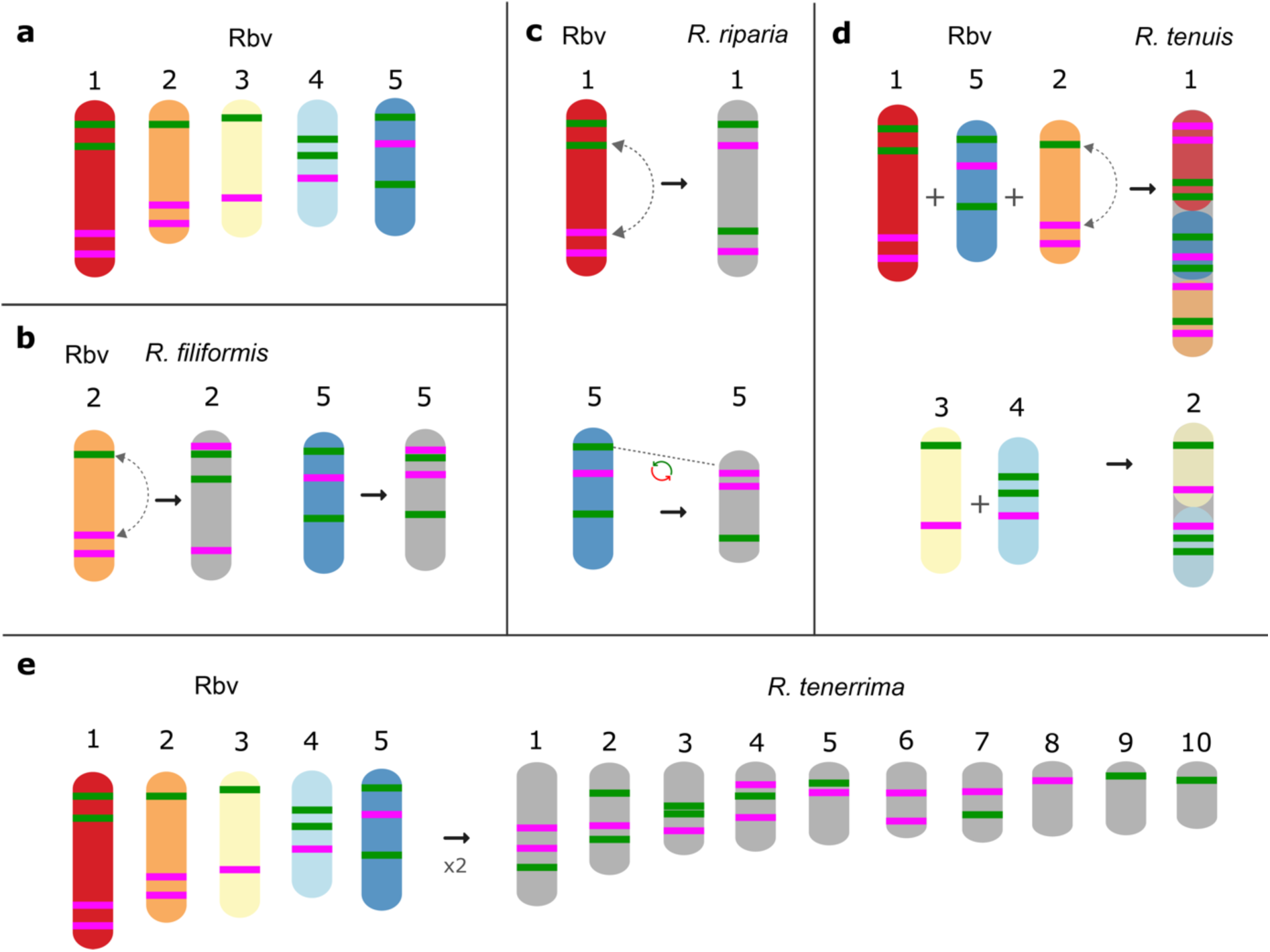
Schematic idiograms of chromosome fusions identified in *Rhynchospora* section *Tenues* species using *R. breviuscula* (a; Rbv) oligo-probes. (**b**) Translocation and additional signal on chromosome 2 and 5 in *R. filiformis.* (**c**) Inversion of the chromosomes 1 and duplication and deletion of magenta and green signals of the chromosomes 5 in *R. riparia.* (**d**) Fusions of the 1+2**+**5 and 3+4 chromosomes of *R. breviuscula* in the two *R. tenuis* chromosomes. (**e**) Barcode signals in *R. tenerrima*, showing a higher number of green (11) and magenta (10) signals compared to the diploid *R. breviuscula*, suggesting this species might be indeed polyploid, but experiencing diploidization.

#### Other clades

In comparison to *R. breviuscula*, the rest of the species analyzed here diverged from a common ancestor ∼35 Mya ago (Costa *et al.,* 2023). The two oligo-probes produced massive background signals on the chromosomes of *R. holoschoenoides* (Rich.) Herter (sect. *Polycephalae* C.B. Clarke) and *R. barbata* (Vahl) Kunth (sect. *Pauciflorae* Kük.; **Fig. 2**. **Fig. 5 & Fig. S1**), suggesting loss of sequence specificity or, less likely, extensive microcollinearity breaks between the genomes. On the other hand, signals were observed on the chromosomes of *R. alba* (*Albae* sect.), *R. cephalotes* (*Cephalotae* sect.) and *R. ridleyi* (*Pseudocapitatae* sect.; **Fig. 2 & Fig. 5**), in congruence with closer phylogenetic distances. However, most of the chromosomes could not be clearly identified based on the barcode pattern observed on *R. breviuscula* chromosomes. The karyotypes of these species have higher chromosome numbers than *R. breviuscula*, up to 13 pairs in *R. alba*. Although the low number of hybridization signals observed for these species could indicate partial sequence divergency, it might also suggest that they have undergone chromosome fissions rather than polyploidy.

**Figure 5:**
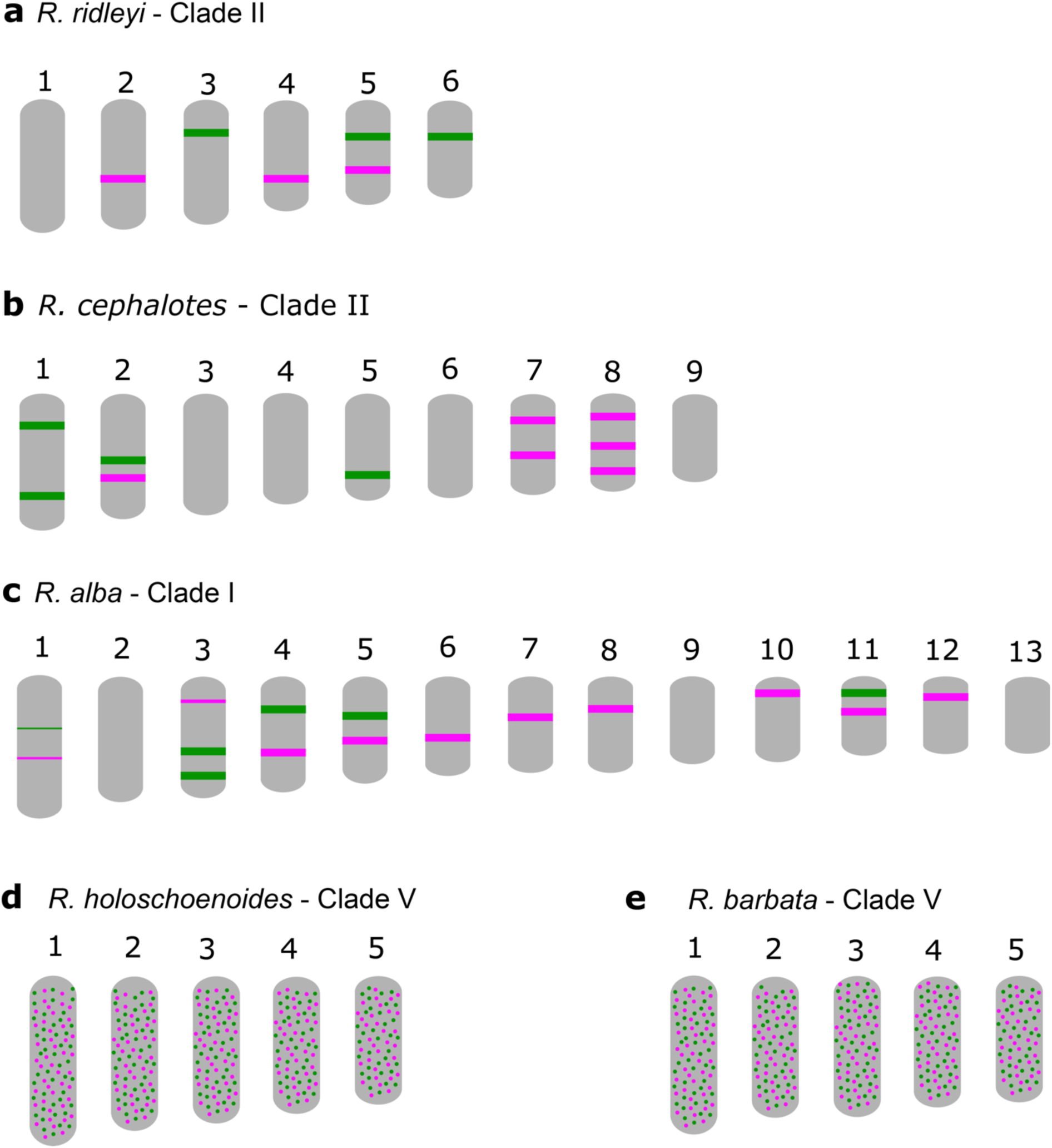
Schematic idiograms of more distant species when compared to *Rhynchospora breviuscula*, showing FISH signals from the Rbv-I (green) and Rbv-II (magenta) oligo-probes. (**a**) Only a few signals were observed in *R. ridleyi*, hampering ortholog identifications. In *R. cephalotes* (**b**) and *R. alba* (**c**), despite the lack of correlation to the *R. breviuscula* barcode pattern, the low number of signals observed suggest that the high chromosome number might have derived from chromosome fissions, rather than polyploidy. In both species of clade V, *R. holoschoenoides* (**d**) and *R. barbata* (**e**) the signals were found dispersed likely, due to lack of specificity, despite sharing the same chromosome number as *R. breviuscula* (*n* = 5).

## DISCUSSION

An optimized oligo-FISH painting method by *de novo* synthesis of thousands of oligos is now possible for species with assembled genomes, providing a powerful tool for understanding the structure, organization, and evolution of plant chromosomes (Braz *et al.,* 2018; Liu *et al.,* 2020; Li *et al.,* 2021; do Vale Martins *et al.,* 2021). Furthermore, oligo-FISH barcoding has also proven to be a powerful and efficient technique for chromosome identification and inference of intrachromosomal rearrangements that are not detected by chromosome painting (Luo *et al.,* 2022; Braz *et al.,* 2018; de Oliveira Bustamante *et al.,* 2021; Nascimento and Pedrosa-Harand, 2023). Here we have successfully developed two sets of oligo-probes based on conserved regions with >20,000 oligos of 45 nt per linkage group, and applied oligo-FISH barcoding to identify all chromosome pairs in *R. breviuscula* karyotype. Among the genus *Rhynchospora*, only three species possess a published genome (Hofstatter *et al.,* 2022) and of those, only *R. breviuscula* is a diploid with 2*n* = 10 chromosomes (reflecting its basic chromosome number *x* = 5; Burchardt *et al.,* 2020), making it the perfect candidate to develop the oligo-probes and transfer them into other species of *Rhynchospora*. We tested the barcode probes in 13 additional species of the genus, which represented seven of the 28 formally described sections (Thomas *et al.,* 2009; Buddenhagen *et al*., 2016; Silva Filho *et al.,* 2021).

In holocentric species, multiple chromosome rearrangements, mainly fusion and fission events, have been shown to be the main driver of their karyotype evolution and may exhibit adaptive potential possibly allowing chromosomal speciation (Hofstatter *et al.,* 2022; Lucek *et al*., 2022; Senaratne *et al.,* 2022; Escudero *et al.,* 2023; Márquez-Corro *et al.,* 2023). In fact, comparisons using recent available chromosome-scale genome assemblies between holocentric *Carex* and *Rhynchospora* species, as well as with their closest monocentric relative *Juncus* L. (rushes) have revealed conserved synteny despite high rates of chromosome fission and fusion in sedges (Hofstatter *et al.,* 2022; Escudero *et al.,* 2023). These data suggest high synteny conservation despite the ancient lineage split between rushes and sedges (>60 Mya) and between *Carex* and *Rhynchospora* (40-50 Mya; Hofstatter *et al*., 2022; Costa *et al.,* 2023). In addition, the recent described chromosome fusions in the genomes of *R. pubera* and *R. tenuis* (Hofstatter *et al*., 2022) could now be confirmed by our oligo barcode probes, showing that this approach is also useful for dynamic holocentric karyotypes. Furthermore, we could detect the chromosomes involved in end-to-end fusions in *R. tenuis* (2*n* = 4), as well as the two rounds of whole genome duplication events and the complex chain of end-to-end fusions that drove the karyotype evolution in *R. pubera* (Hofstatter *et al.,* 2022).

Our oligo barcode analysis further allowed us to better understand an ongoing karyotype differentiation among different populations of *R. pubera*. Our reference *R. pubera* (2*n* = 10) from the Brazilian northeast (Hofstatter *et al*., 2022) is the most common cytotype found for this species, but earlier studies have reported a population from Manaus, in the Amazonas state in North Brazil, with 2*n* = 12 (Arguelho *et al*., 2012). Here, we report a newly collected sample from another population from North Brazil, from Belém in the Pará state, also showing 2*n* = 12. Here, we report a newly collected sample from another population from North Brazil from Belém in the Pará state showing also 2*n* = 12. The oligo barcode pattern observed in *R. pubera*-12 indicate that its chromosomes originated from the same duplication and fusions events as observed in *R. pubera*-10, but also suggest ongoing complex chromosome genome reshuffling, leading to the ascending dysploidy, in a likely scenario of ongoing chromosomal speciation. Apparently, in contrary to the conservation of all four copies in the “standard” *R. pubera* karyotype, this accession has lost sequences corresponding to approximate one whole set of ancestral chromosomes. Such complex rearrangements are supported by the meiotic irregularities previously reported for a *R. pubera* cytotype with 2*n* = 12 (Arguelho et al., 2012). However, the limitations of the barcode probes, particularly when some signals were lost during genome reshuffling, underscore the need for a comprehensive genomic study to validate these findings.

Dual-color oligo-FISH is useful for studying species evolution and previous research has shown that oligo probes can be applied to chromosome identification in related species that diverged ∼5 My ago, such as *Phaseolus* (Nascimento and Pedrosa-Harand, 2023), ∼12 Myr as in *Cucumis* (Hans *et al.,* 2015) or *Sorghum* (Yu *et al.,* 2021), ∼15 Mya as in *Solanum* species (Braz *et al.,* 2018), or even up to ∼18 Mya as in *Saccharum* (Yu *et al.,* 2021). Here we transferred *R. breviuscula* oligo-probes and demonstrated their utility for correct ortholog identification among species with divergence times up to ∼25 My (clade IV, sections *Dichromena* and *Tenues*). Indeed, all true diploid species of sect. *Dichromena* with *n* = 5 (*R. colorata*, *R. nervosa* and *R. radicans*) showed exactly the same barcode pattern as *R. breviuscula*. Similarly, species of the related sect. *Tenues* showed a very similar pattern, but with the occurrence of several recognizable rearrangements, e.g., fusions, inversions and translocations. Additionally, we could also detect a potential polyploidy event in *R. tenerrima*. However, the reduced number of signals observed in contrast to the expected duplicated amount of barcode signals suggest that this species is undergoing rediploidization. Indeed, no meiotic irregularities was reported for *R. tenerrima*, where a normal pollen development was observed (Arguelho *et al*., 2012). Low sequence similarity or the interruption of synteny have hampered the barcode recognition between species with divergence time up to ∼30 My from *R. breviuscula*.

In general, our results indicate a good relationship between chromosomal rearrangements with the current understanding of the phylogenetic relationships in *Rhynchospora*. Our oligo barcode results confirmed, as observed in other sedges, that: (i) there is high synteny among holocentric genomes during long periods of time; (ii) that in the genus *Rhynchospora*, the ancestral karyotypes with *x* = 5 chromosomes may be either conserved or a result of fusions and genomic reshuffle events; and (iii) that chromosome rearrangements after fission events appear to govern karyotypic diversity in the genus. The stasis of diploid karyotypes with 2*n* = 10 contrasts with the multiple rearrangements associated to, but not directly involved, in the dysploidy events, as also observed in monocentrics (Nascimento and Pedrosa-Harand, 2023). Ultimately, the current oligo-probe sets enabled the direct visualization of homologous regions on *Rhynchospora* chromosomes through a straightforward experimental method. These innovative oligo-barcode-based comparative FISH studies offer a potent instrument for illuminating the evolutionary dynamics of holocentric beaksedge genomes. This toolset can be readily employed to pinpoint additional *Rhynchospora* karyotypes resulting from chromosome fusion, fission, or polyploidy events, thereby unveiling potential chromosomal reorganization within evolutionarily divergent genomes.

## ACKNOWLEDGMENTS

We acknowledge the excellent technical assistance of Christina Philipp and Ursula Pfordt for keeping the plants growing in the greenhouse at the MPIPZ. This work was conducted during a PhD sandwich fellowship awarded to YMS and LMP supported by the International Cooperation Program PROBRAL from CAPES – Brazilian Federal Agency for Support and Evaluation of Graduate Education within the Ministry of Education of Brazil. This work was supported by a grant awarded to A.P.H (PROBRAL CAPES/DAAD project number 88881.144086/2017-01). G.S. receive productivity fellowship from CNPq (process numbers PQ-312852/2021-5). AM thanks to the Max Planck Society and DFG (grant number MA 9363/3-1) for financial support. We thank Prof. Julio Cesar Pieczarka from Federal University of Pará for collecting and providing us the *R. pubera* (2*n* = 12) sample. We thank the administration of the Biosphere Reserve “*Oberlausitzer Heide- und Teichlandschaft*” for providing occurrence information and sampling permit for *R. alba*. Sample permits for newly collected Brazilian samples (*R. cephalotes*, *R. barbata*, *R. filiformis*, *R. holoschoenoides*, *R. pubera*-12, *R. ridleyi* and *R. tenerrima*) were obtained from Sisgen (registration numbers AF0D72D, A13DD4F, AA62E92). The sample for *R. radicans* from Costa Rica was obtained in collaboration with Andrés Gatica-Aria under a Material Transfer Agreement.

## AUTHOR CONTRIBUTIONS

**Yennifer Mata-Sucre:** Formal analysis, investigation, data curation, writing – original draft, writing – review & editing, visualization. **Letícia Maria Parteka:** Formal analysis, investigation, data curation. **Christiane Ritz, Wayt Thomas, Leonardo P. Felix, Andrés Gatica, André L. L. Vanzela:** Plant material collection and identification – review & editing. **Gustavo Souza, André L. L. Vanzela:** Supervision and visualization – review & editing. **Andrea Pedrosa-Harand, André Marques:** Conceptualization, supervision, resources, writing – review & editing, funding acquisition.

## DATA AVAILABILITY

All data generated or analyzed during this study are included in the manuscript or as supplementary materials.

## MATERIALS AND METHODS

### Plant material

Samples from *R. breviuscula*, *R. cephalotes*, *R. nervosa* ssp. *ciliata*, *R. pubera* (2*n* = 10) and *R. tenuis* were already available from previous studies (Hofstatter *et al*., 2022; Ribeiro *et al*., 2016; Costa *et al*., 2021). The *R. colorata* (L.) H.Pfeiff. sample was commercially acquired. Additional 9 samples from 9 species were collected from different localities in Brazil, Costa Rica and Germany (**Table S3**) and further cultivated under controlled greenhouse conditions (16h daylight, 26 °C, >70% humidity) at the Max Planck Institute for Plant Breading Research in Germany. Sampling includes *Rhynchospora alba* (L.) Vahl (Germany, Sachsen, Kreba-Neudorf, Weißes Lug, DD: 51.355055 N; 14.729016 E), *R. barbata* (Vahl) Kunth (Areia-PB, Brazil), *R. filiformis* Vahl (Baía da Traição-PB, Brazil), *R. holoschoenoides* (Rich.) Herter (Baía da Traição-PB, Brazil), *R. pubera* (Vahl) Boeckeler accession with 2*n* = 12 (Belém-PA, Brazil), *R. radicans* (Schltdl. & Cham.) H.Pfeiff. (San José, Costa Rica), *R. ridleyi* C.B.Clarke (Areia-PB, Brazil), *R. riparia* (Nees) Boeckeler (Rio Tinto-PB, Brazil), *R. tenerrima* Nees ex Spreng. (Vale do Codó-PR, Brazil) and *R. tenuis* Link (Ipojuca-PE, Brazil). These taxa represent species in the sections *Albae*, *Dichromena*, *Cephalotae*, *Pauciflorae, Polycephalae*, *Pseudocapitatae* and *Tenues* (Thomas *et al.,* 2009; Buddenhagen *et al*., 2016; Silva Filho *et al.,* 2021).

### Oligo probe design

Oligo-FISH barcode probes were designed against the chromosome-scale sequence assembly of *R. breviuscula* (Hofstatter *et al*., 2022) using Daicel Arbor Biosciences’ proprietary software (Daicel Arbor Bioscience, Ann Arbor, MI, USA). Briefly, target sequences were fragmented into 43–47 nucleotide-long overlapping probe candidate sequences that were compared to the rest of the genome sequence to exclude any candidates with potential cross-hybridization based on a predicted Tm of hybridization. Non-overlapping target-specific oligonucleotides were selected for the final probe sets and synthesized as myTAGs® Labeled Libraries (Daicel Arbor Bioscience). Target regions were selected with a relatively high density of oligos based on the density distribution profile on the entire chromosome (**Table S1**), and then taking into account the regions conserved with *R. pubera* and *R. tenuis* chromosome-scale sequence assemblies (Hofstatter *et al*., 2022). The two libraries consist of ∼ 20,000 oligomers each (45nt long) corresponding to eight green (Rbv-I) and seven magenta (Rbv-II) signals. Each chromosome of *R. breviuscula* has two to four unique signals, with specific combinations of green and magenta signals covering specific chromosomes regions (**Fig. 1**). The libraries were synthesized by Arbor Biosciences (Ann Arbor, Michigan, USA) and directly labeled with Cy3 and Alexa-488 (Eurofins Genomics, Ebersberg, Germany).

### Chromosome spreads and oligo-Fluorescence *in situ* hybridization (Oligo-FISH)

To prepare mitotic metaphase chromosomes, root tips were fixed in ethanol:acetic acid (3:1 v/v) for 2–24 h at room temperature and stored at -20 °C. Fixed root tips were washed twice in distilled water and digested in an enzymatic solution of 2% (w/v) cellulase (Onozuka)/20% (v/v) pectinase (Sigma) at 37 °C, for 90 min. Meristems were macerated in a drop of 45% acetic acid and spread on a hot plate following Ruban *et al*. (2014). Oligo-FISH was performed according to the protocol proposed by Braz *et al*. (2020) with some modifications. The hybridization mixture consisted of 50% formamide, 2×SSC (sodium citrate saline; pH 7.0), 10% dextran, 350 ng of Alexa-labeled probe and 200 ng of Cy3-labeled probe, in a total volume of 10 µl per slide. Chromosomes were denatured for 5-7 min at 75°C and incubated for 18-24 h at 37 °C in a humid chamber. Subsequently, coverslips were gently removed and the slides were washed with 2× and 0.1× SSC at 42 °C (∼ 76% final stringency). In addition, low-stringency baths were also performed in more distant species, following Braz *et al*. (2020). Chromosomes were counterstained with 2 µg/mL DAPI in Vectashield solution (Vector Laboratories).

### Image processing and comparative analyses

Images of the chromosomes were captured with a Zeiss Axiovert 200M microscope equipped with a Zeiss AxioCam CCD. Images were analyzed using the ZEN software (Carl Zeiss GmbH, Jena Germany). Karyograms were obtained from the best cell photographed using Adobe Photoshop CS5 (version 12.0) and ordered following their orthologies. Chromosome size measurements of the 14 species studied (Table S2) were constructed after analysis of at least five whole metaphase cells using DRAWID 0.26 software (Kirov *et al*., 2017) and interpreted using the genetic relationships described in Costa *et al*. (2023).

## Supplementary tables

**Table S1.** Oligo-barcode design showing the distribution of 45nt-oligo density across the five chromosomes of *R. breviuscula*.

**Table S2.** Chromosome size in 14 *Rhynchospora* species.

**Table S3.** Sample information, voucher and geographical coordinates.

## Supplementary figures

**Figure S1:**
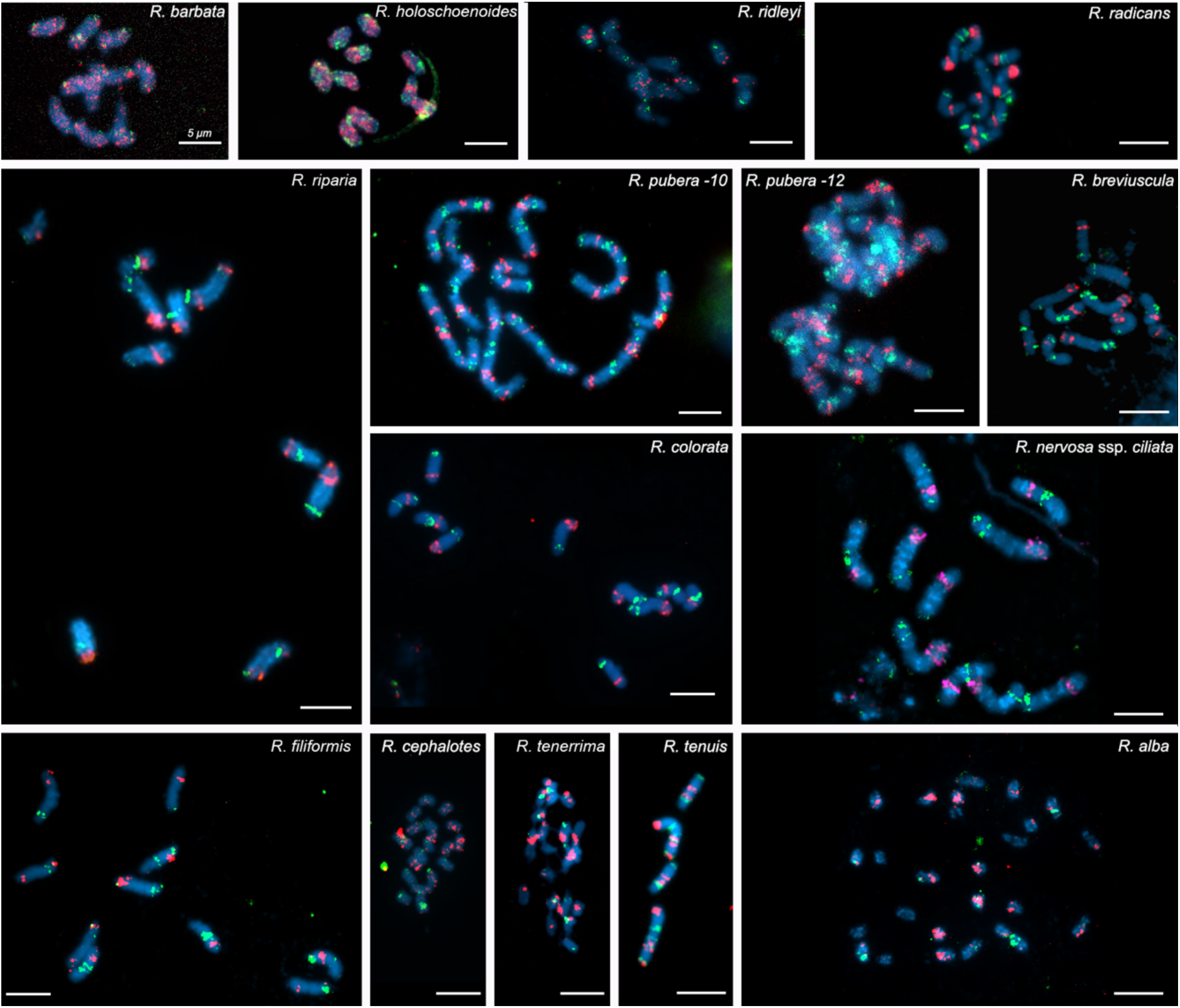
Chromosome identification by oligo-FISH barcoding in *Rhynchospora* species using Rbv-I (green) and Rbv-II (magenta) oligo probes developed based on *R. breviuscula* (*n* = 5) reference genome. *R. barbata* 2*n* = 10, *R. holoschoenoides* 2*n* = 10, *R. ridleyi* 2*n* = 12, *R. radicans* 2*n* = 10, *R. riparia* 2*n* = 10, *R. pubera* 2*n* = 10, *R. pubera* 2*n* = 12, *R. breviuscula* 2*n* = 10, *R. colorata* 2*n* = 10, *R. nervosa* subsp. *ciliata* 2*n* = 10, *R. filiformis* 2*n* = 10, *R. cephalotes* 2*n* = 18, *R. tenerrima* 2*n* = 20, *R. tenuis* 2*n* = 4, and *R. alba* 2*n* = 26. Note that despite some similar patterns to those of *R. breviuscula*, we could not decipher the complete karyotype of *R. pubera*-12. Bars = 5 µm.

**Figure S2:**
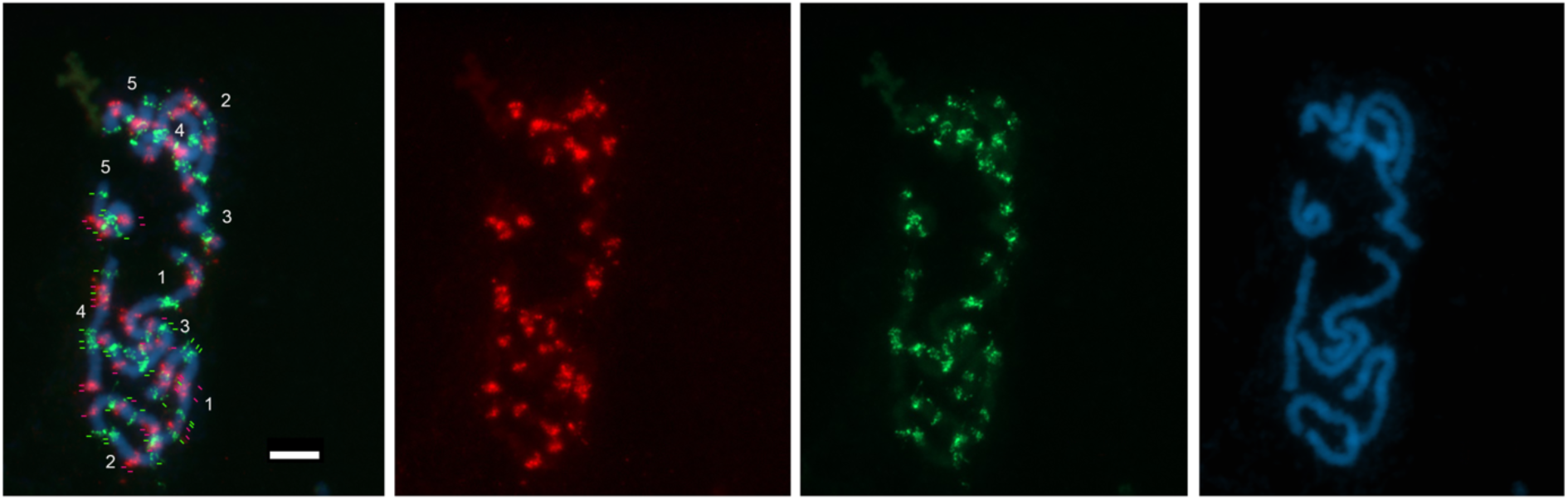
Chromosome identification in prometaphase cells by oligo-FISH barcoding in *R. pubera*-10 using Rbv-I (green) and Rbv-II (magenta) probes developed in *R. breviuscula*. 8×4 green and 7×4 magenta signals are shown with a total of 60 signals in the haploid genome of *R. pubera*-10. Bar = 5 µm.

